# New oligodendrocytes exhibit more abundant and accurate myelin regeneration than those that survive demyelination

**DOI:** 10.1101/2020.05.22.110551

**Authors:** Sarah A Neely, Jill M Williamson, Anna Klingseisen, Lida Zoupi, Jason J Early, Anna Williams, David A Lyons

## Abstract

Regeneration of myelin (remyelination) in the central nervous system (CNS) has long been thought to be principally mediated by newly generated oligodendrocytes, a premise underpinning therapeutic strategies for demyelinating diseases, including multiple sclerosis (MS). Recent studies have indicated that oligodendrocytes that survive demyelination can also contribute to remyelination, including in MS, but it is unclear how remyelination by surviving oligodendrocytes compares to that of newly generated oligodendrocytes. Here we studied oligodendrocytes in MS, and also imaged remyelination *in vivo* by surviving and new oligodendrocytes using zebrafish. We define a previously unappreciated pathology in MS, myelination of neuronal cell bodies, which is recapitulated during remyelination by surviving oligodendrocytes in zebrafish. Live imaging also revealed that surviving oligodendrocytes make very few new sheaths, but can support sheath growth along axons. In comparison, newly made oligodendrocytes make abundant new sheaths, properly targeted to axons, and exhibit a much greater capacity for regeneration.

## Introduction

Damage to, or loss of myelin (demyelination) is a characteristic feature of many debilitating disorders of the CNS, including MS (1, 2). However, the CNS can regenerate myelin, and promoting remyelination represents a therapeutic goal due to the hope that this will help restore lost function and protect demyelinated axons (3). In various animal models, demyelination is followed by a robust activation of oligodendrocyte progenitor cells (OPCs), which proliferate and migrate to sites of damage, where they can generate new oligodendrocytes that remyelinate axons (1). In addition, there is evidence from pathological and imaging studies that remyelination can occur in MS, but it is limited, variable, imperfect, and myelin loss accumulates over time (2, 4, 5). Although the mechanisms that facilitate or inhibit remyelination remain to be fully elucidated, OPCs have been observed in many demyelinated lesions, suggesting that their failure to generate mature oligodendrocytes represents a bottleneck to regeneration (3, 6). This premise has led to the search for and identification of drug candidates that can promote oligodendrocyte generation and remyelination (7–9), several of which have progressed to clinical trial (10). Although the results of these trials have been mixed to date, there are indications that this strategy may provide clinical benefit (11). More recent studies have indicated diversity in oligodendrocytes in disease (12), including the possibility that oligodendrocytes that survive demyelination contribute to remyelination in MS (13). Remyelination by oligodendrocytes that survive demyelination has now also been documented in animal models (14, 15), which, if possible to promote, could represent an additional opportunity to enhance regeneration.

Despite the greatly different environment that follows demyelination, OPCs redeploy largely the same strategies to generate oligodendrocytes during remyelination as they do in the healthy nervous system (16), and we have a comprehensive understanding of the mechanisms underpinning oligodendrocyte differentiation (17). In contrast, insights into how oligodendrocytes interact with axons to coordinate the process of myelination per se are only emerging (18), and thus our understanding of the cell biology of remyelination remains limited. In the healthy nervous system, individual oligodendrocytes extend numerous exploratory processes that interact with many prospective targets, making myelin sheaths on certain axons, while avoiding others, a process that occurs over the order of hours (19–22). After their formation, the growth and remodelling of sheaths occurs slowly over a more protracted phase of days to weeks, and albeit more limited in extent, throughout life (23–26). Therefore, one might predict that new oligodendrocytes undergoing remyelination would also exhibit an initial highly dynamic restricted period of target selection and sheath formation, followed by a slower protracted period of sheath growth and remodelling. A restricted period of sheath formation begs the question as to how oligodendrocytes that survive demyelination contribute to remyelination. Can oligodendrocytes that survive demyelination extend new dynamic processes that interact with prospective targets? If so, do they form a large number of processes and correctly target sheaths that grow to relatively normal lengths? If not, might surviving oligodendrocytes exhibit very limited, slow, or even aberrant remyelination? Given that mature myelin sheaths can renew growth (27), and be stimulated to produce more myelin during remyelination (28), might the regenerative contribution of surviving oligodendrocytes be limited to supporting renewed growth of sheaths that have not been completely lost? To answer these questions requires models in which both surviving and new oligodendrocytes can be followed over time during remyelination.

The observation that surviving oligodendrocytes can make new myelin sheaths has only recently been documented (14). The fact that this has not been noted before may reflect the fact that oligodendrocyte death is closely associated with demyelination in most standard models of demyelination, leaving limited opportunity to study surviving cells. In contrast, studies in MS have indicated diversity in modes of demyelination, oligodendrocyte pathology and state, including myelin loss without cell death (12, 29, 30). Therefore, we aimed to combine analyses of oligodendrocytes in MS with longitudinal observations of remyelination using a new zebrafish model that allowed us to compare the cell biology of remyelination by surviving and new oligodendrocytes.

## Results

### Myelin is mistargeted to neuronal cell bodies during remyelination in MS

To better understand oligodendrocytes in MS we analysed human brain post-mortem motor cortex samples from 5 people with, and 5 people without MS (**Figure 1**). We focused our analyses of oligodendrocytes and myelin on the grey matter because the lower density of myelin in this area allows discernment of individual oligodendrocytes and discrete myelin sheaths. We first compared PLP-positive myelin sheaths within demyelinated lesions, perilesional areas and adjacent normal appearing grey matter. This revealed numerous aberrant-appearing profiles with PLP-labelling appearing to enwrap cells, suggestive of myelin mistargeting. Therefore, we next co-stained MS tissue with PLP and NeuN (to label neuronal cell bodies) and found numerous PLP-labelled structures enwrapping NeuN-labelled neuronal cell bodies (**Figure 1A-E**). Quantification showed that the number of myelinated neuronal cell bodies in MS perilesion sites, where remyelination is thought to take place (31), was almost 100-fold higher compared to control (non-MS) grey matter or normal appearing grey matter in MS (**Figure 1F, Figure S1**). These data indicate that myelin mistargeting is a previously unappreciated feature of MS grey matter pathology. Given the localisation of oligodendrocytes exhibiting myelin mistargeting to areas of prospective remyelination, we wondered whether this pathology might reflect impaired remyelination, either by newly generated oligodendrocytes or oligodendrocytes that survive demyelination that have recently been proposed to be common in MS (13). To test this possibility, and directly compare remyelination by both surviving and newly generated oligodendrocytes, we developed a novel demyelination model using zebrafish.

**Fig. 1.**
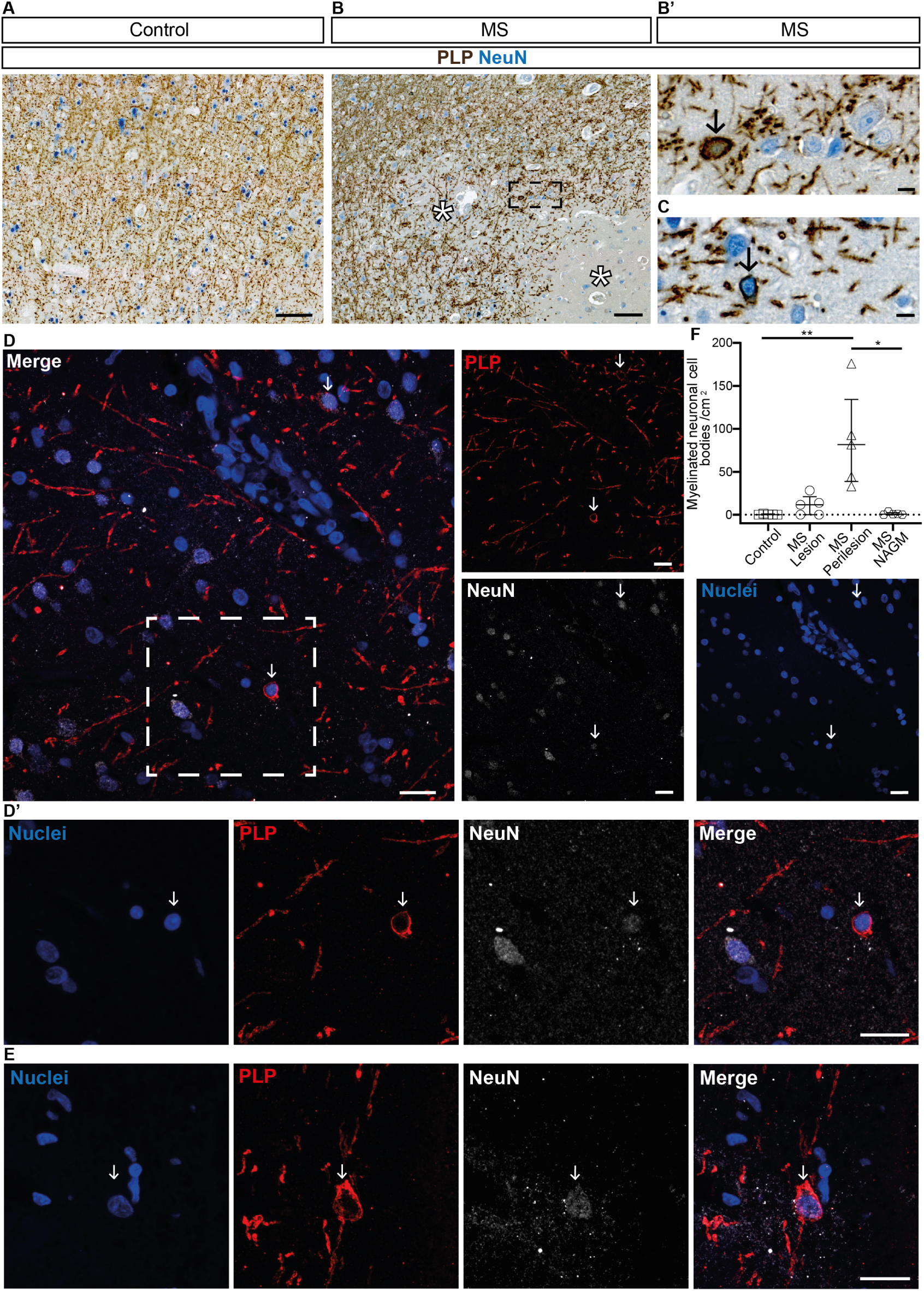
Mistargeted myelin profiles are present in remyelinating lesions in motor cortex tissue from people with MS. (A and B) Low magnification images of chromogenic immunohistochemistry for proteolipid protein (PLP - brown) and NeuN (blue) in (A) human control (people without MS) motor cortex and (B) human MS motor cortex. Asterisks indicate lesion areas. Scale bars, 100 µm. (B’ and C) High magnification images of myelin-wrapped (PLP - brown) neuronal cell bodies (NeuN - blue) in human MS motor cortex, at positions indicated by black arrows. (B’) shows a high magnification view of the region highlighted in the black box in (B). Scale bars, 10 µm. (D) Low magnification image of fluorescent immunohistochemistry for NeuN (white), PLP (red) and Hoechst (nuclei-blue) in human MS motor cortex. White arrows indicate locations of myelin wrapped (PLP +ve) neuronal (NeuN +ve) cell bodies (Hoechst +ve). Merged channel image scale bar, 20 µm. Split channel scale bars, 10 µm. (D’ and E) High magnification images of fluorescent immunohistochemistry for NeuN (white), PLP (red) and Hoechst (nuclei-blue) in human MS motor cortex. White arrows indicate the location of myelin wrapped (PLP +ve) neuronal (NeuN +ve) cell bodies (Hoechst +ve). (D’) shows a high magnification view of the region highlighted in the white box in (D). Scale bars, 20 µm. (F) Quantification of the number of myelinated neuronal cell bodies per grey matter area (cm^2^) in control grey matter (median = 0.00, 25^th^ percentile 0.00, 75^th^ percentile 0.82), MS lesion area (median = 11.65, 25^th^ percentile 0.00, 75^th^ percentile 21.06), MS perilesion area (median = 81.63, 25^th^ percentile 38.65, 75th percentile 134.30) and MS normal appearing grey matter (NAGM) (median = 0.61, 25^th^ percentile 0.00, 75^th^ percentile 2.32). Control versus MS lesion p > 0.9999, control versus MS perilesion p = 0.0076, control versus MS NAGM p > 0.9999, MS lesion versus MS perilesion p = 0.1735, MS lesion versus MS NAGM p > 0.9999, MS perilesion versus MS NAGM p = 0.0321. Kruskal-Wallis test with Dunn’s multiple comparisons test. All samples are from human motor cortex tissue: N = 5 human tissue samples (from 5 different individuals) per condition.

### A zebrafish model to study remyelination by both oligodendrocytes that survive demyelination and newly generated oligodendrocytes

Given their advantages for longitudinal live imaging of myelination at high resolution *in vivo* (32), we designed a zebrafish model to follow the fate of single oligodendrocytes after demyelination. We wanted to generate a model in which we could induce extensive demyelination, but where a significant number of affected oligodendrocytes might survive. We reasoned that stimulating a large and localized influx of cations including Ca^2+^ into myelin sheaths might induce demyelination, without necessarily leading to cell death. This was based on our observation that high amplitude, long duration Ca^2+^ transients prefigure myelin sheath retraction during development (19), and that excitotoxic influx of cations can mimic myelin pathology in mammals (33, 34). To induce cation influx into myelinating oligodendrocytes we made a transgenic zebrafish line, Tg(mbp:TRPV1-tagRFPt), in which we expressed the rat form of the capsaicin (csn)-inducible, cationpermeable transient receptor potential V1 channel (TRPV1) in myelinating glia (**Figure 2A and Methods**). Importantly, unlike the rat form of the protein, zebrafish TRPV1 channels are not csn sensitive (35), and treatment of zebrafish with csn alone had no observable adverse effects on myelin or oligodendrocytes (**Figure S2C**). To induce myelin damage we treated Tg(mbp:TRPV1-tagRFPt) animals with 10 µM csn for 2 hours at 4 days post fertilisation (dpf) (**Figure S2 and Methods**), a time by which many axons of the larval zebrafish spinal cord are myelinated (36). This led to extensive demyelination, as assessed by the transgenic reporter Tg(mbp:EGFP-CAAX) as well as transmission electron microscopy (TEM) (**Figure 2B and I-L**). Indeed, our TEM analyses revealed a 95 % reduction in myelinated axon number in the dorsal tract of the spinal cord just 1 day post treatment (dpt) with a concomitant increase in the number of unmyelinated axons, indicating no axonal loss at this point (**Figure 2I-L**). Importantly, despite this extensive loss of myelin, we saw a relatively modest reduction in total oligodendrocyte number, with 33 % fewer oligodendrocytes in csn treated Tg(mbp:TRPV1-tagRFPt) animals at 3 hpt (**Figure 2C-H**). Following this, oligodendrocyte number increased in csn-treated animals and also controls, with numbers becoming indistinguishable by 5 dpt (**Figure 2C-H**). The oligodendrocytes generated (**Figure 2C**), and the myelin that they produce (**Figure 2B**) during this time were located in the same regions in control and csn-treated animals, indicating that remyelination of demyelinated axons takes place in csn-treated animals. Together, these observations indicate that this model allows investigation of remyelination by both oligodendrocytes that survive demyelination and those newly generated following demyelination.

**Fig. 2.**
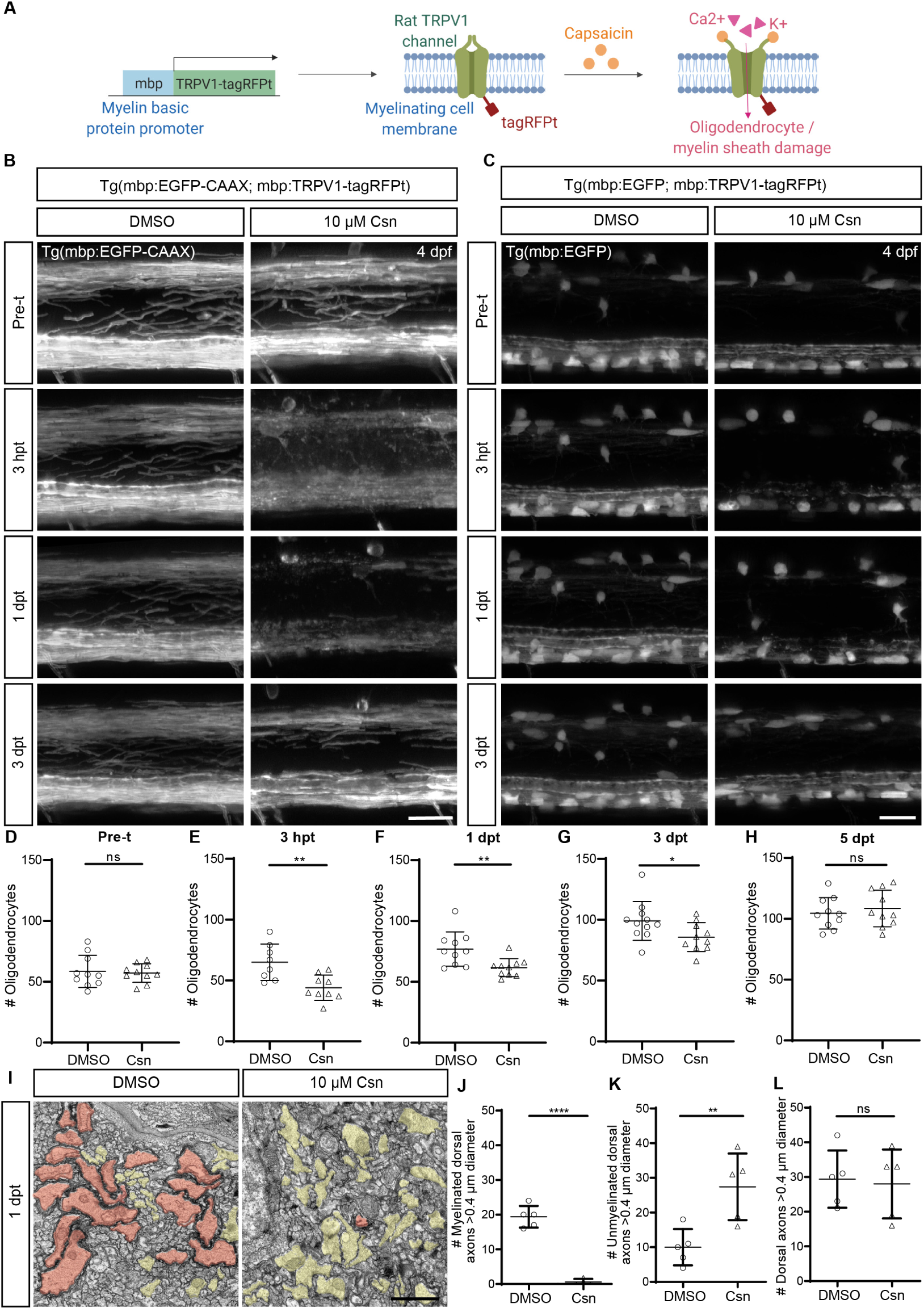
Demyelination with minimal oligodendrocyte loss in the Tg(mbp:TRPV1-tagRFPt) demyelination model in zebrafish. (A) Schematic illustrating the Tg(mbp:TRPV1-tagRFPt) demyelination model. The rat form of the TRPV1 channel is expressed in myelinating oligodendrocytes and is activated by addition of csn which drives cation influx. Zebrafish TRPV1 channels are insensitive to csn, therefore csn treatment specifically results in damage to myelinating glia which express the rat form of the TRPV1 channel. (B and C) Confocal images of (B) myelin visualised in Tg(mbp:EGFP-CAAX; mbp:TRPV1-tagRFPt) zebrafish and (C) myelinating oligodendrocytes visualised in Tg(mbp:EGFP; mbp:TRPV1-tagRFPt) zebrafish, with control (DMSO) and csn treated animals in left and right panels respectively pre-treatment (pre-t) at 4 dpf, 3 hpt, 1 dpt and 3 dpt. Scale bars, 20 µm. (D-H) Quantification of myelinating oligodendrocyte number in DMSO and csn treated Tg(mbp:EGFP; mbp:TRPV1-tagRFPt) zebrafish over time. (D) Pre-treatment DMSO (mean = 58.60 ± 13.23 SD) versus csn (mean = 57.20 ± 7.642 SD), p = 0.7753. (E) 3 hpt DMSO (mean = 65.00 ± 14.93 SD) versus csn (mean = 44.11 ± 10.36 SD), p = 0.0041. (F) 1 dpt DMSO (mean = 76.90 ± 14.14 SD) versus csn (mean = 61.70 ± 7.39 SD), p = 0.0075. (G) 3 dpt DMSO (mean = 99.00 ± 15.93 SD) versus csn (mean = 85.70 ± 11.83 SD), p = 0.0444. (H) 5 dpt DMSO (mean = 104.5 ± 12.95 SD) versus csn (mean = 108.5 ± 15.10 SD), p = 0.5328. Unpaired two-tailed t-test. N = 8 - 11 zebrafish per condition. Each data point represents total (dorsal + ventral) oligodendrocyte number analysed per imaged area per zebrafish. (I) Transmission electron microscopy images of DMSO and csn treated Tg(mbp:TRPV1-tagRFPt) zebrafish at 1 dpt show numerous large calibre myelinated axons (red) in DMSO and numerous large calibre unmyelinated axons (yellow) in csn treated animals. Scale bar, 1 µm. (J) Quantification of the number of myelinated axons in the dorsal spinal cord at 1 dpt in DMSO (mean = 19.40 ± 3.13 SD) versus csn (mean = 0.600 ± 0.89 SD), p < 0.0001. Unpaired two-tailed t-test with Welch’s correction. N = 5 zebrafish per condition. (K) Number of unmyelinated axons in the dorsal spinal cord at 1 dpt in DMSO (mean = 10.00 ± 5.24 SD) versus csn (mean = 27.40 ± 9.61 SD), p = 0.0075. Unpaired two-tailed t-test. N = 5 zebrafish per condition. (L) Number of large calibre axons (diameter > 0.4 µm) at 1 dpt in DMSO (mean = 29.40 ± 8.26 SD) versus csn (mean = 28.00 ± 9.95 SD), p = 0.8148. Unpaired two-tailed t-test. N = 5 zebrafish per condition.

### Oligodendrocytes that survive demyelination exhibit limited new myelin sheath generation and mistarget their newly made myelin

To follow the fate of individual oligodendrocytes (both surviving and newly generated cells), we mosaically labelled single cells by injection of mbp:EGFP-CAAX plasmid DNA (**Methods**). To follow the fate of oligodendrocytes that survive demyelination, we imaged oligodendrocytes prior to the induction of demyelination, 3 hours post csn treatment (hpt), 1 dpt, and 3 dpt (**Figure 3**). This confirmed that oligodendrocytes could survive complete loss of their myelin sheaths (**Methods, Figure S2**). Of the 37 oligodendrocytes analysed in 37 animals from prior to demyelination, 23 oligodendrocytes survived for at least 3 dpt. Of these surviving oligodendrocytes, 22 cells made new myelin during this time (**Figure 3B-C, and E**), indicating that oligodendrocytes either make new myelin or undergo cell death. Detailed morphological analysis of these oligodendrocytes allowed us to assess several parameters including myelin sheath number and length as well as the presence of any aberrant myelin profiles. Although almost all surviving cells made new myelin, these oligodendrocytes made very few new sheaths, producing 2 sheaths per cell on average during remyelination, compared with the average of 18 that they had prior to demyelination (**Figure 3G**). One of the most striking observations we made was that the new myelin that these cells made was often mistargeted. Of the 22 surviving oligodendrocytes that made new myelin, the majority (13/22, 59 %) mistargeted myelin, as evidenced by the appearance of circular myelin profiles surrounding cell bodies (**Figure 3B, F**), as we have reported in other contexts (37, 38), and similar to that observed in our analyses of MS tissue. In contrast, only 1 of these same oligodendrocytes exhibited any evidence of myelin mistargeting prior to demyelination (**Figure 3F**). This suggests that extensive myelin mistargeting is a pathological feature of remyelination by oligodendrocytes that survive demyelination. We also observed that new sheaths and aberrant mistargeted myelin profiles were localised close to surviving oligodendrocyte cell bodies (**Figure 3B+C**), indicating that these cells have very restricted process dynamics. Together our observations indicate that mature oligodendrocytes have a limited capacity to reinitiate dynamic process activity, new sheath formation, or accurate myelin targeting. Despite the very limited formation of new sheaths by cells that survived demyelination, we found that when sheaths were occasionally made on axons, there was no significant difference in their average length compared to those made by the same cells prior to demyelination (**Figure 3H**). This suggests that surviving oligodendrocytes do retain a capacity to support renewed growth of sheaths along axons, and that the bottleneck to successful remyelination is in the earlier phase of processes extension, targeting and sheath formation. Therefore, we wanted to test whether enhancing their process extension and ability to make new sheaths improved their remyelination capacity. To do so, we inhibited Rho Kinase (ROCK) following demyelination, using the inhibitor Y27632, which increases the number of myelin sheaths made by oligodendrocytes in zebrafish and mammals (39–41). We found that inhibition of ROCK did not significantly affect the number or length of myelin sheaths made on axons by oligodendrocytes that survived demyelination, but instead increased yet further the mistargeting of myelin by these cells (**Figure 3D and I-J**). Collectively our data indicate that oligodendrocytes that survive demyelination have a poor capacity for accurate reformation of new myelin sheaths *in vivo*.

**Fig. 3.**
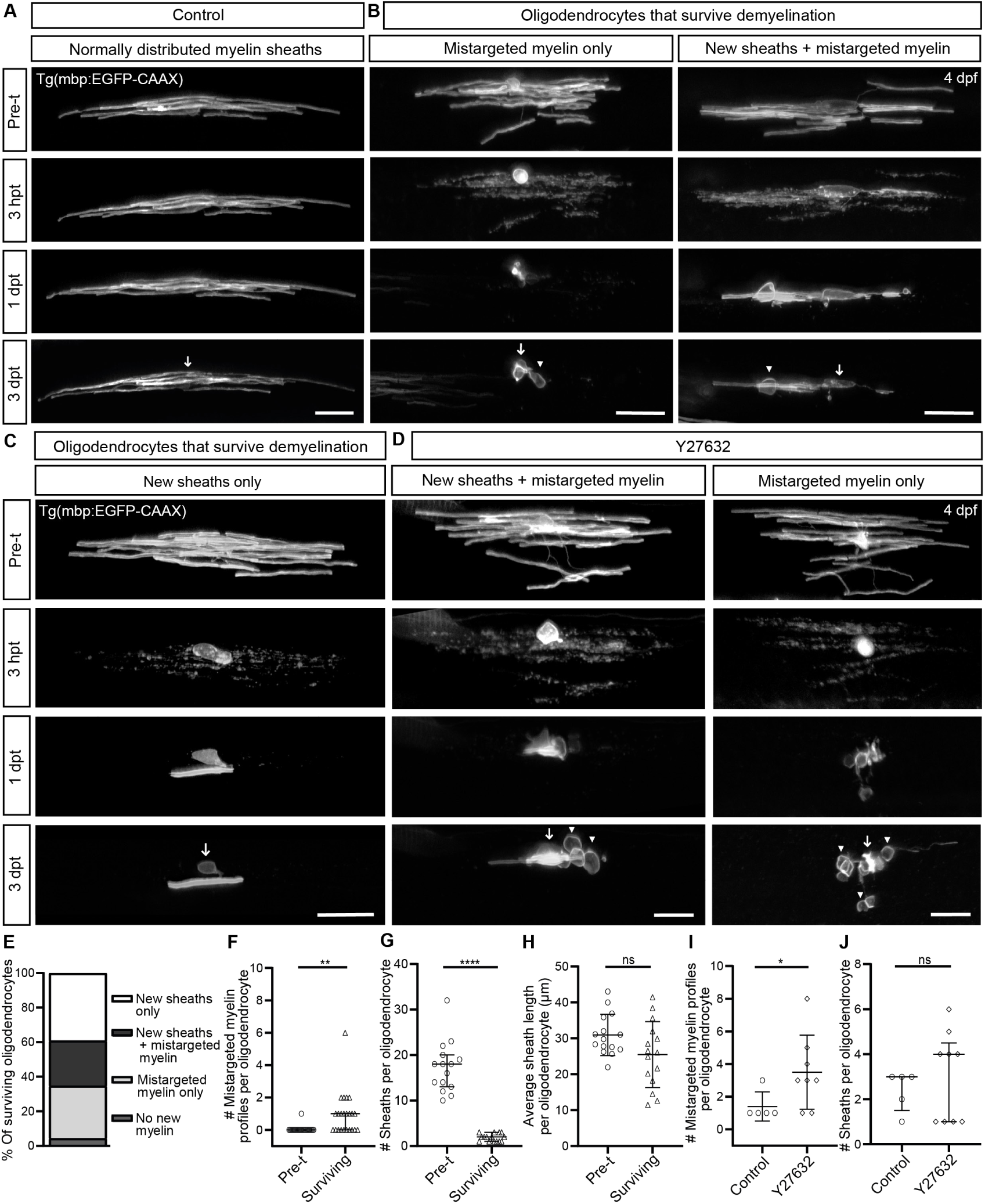
Remyelination by oligodendrocytes that survive demyelination. (A-C) Confocal images of single oligodendrocytes followed over time from pre-treatment (pre-t) to 3 dpt. Arrows show position of cell bodies. Arrowheads show positions of mistargeted myelin. (A) Image of a single oligodendrocyte in a control (DMSO) treated zebrafish. (B) Images of single oligodendrocytes in csn treated zebrafish which mistarget their newly made myelin. (C) Image of a single oligodendrocyte in a csn treated zebrafish which targets its newly made myelin correctly. Scale bars, 20 µm. (D) Confocal images of oligodendrocytes followed over time from prior to csn treatment through to 3 dpt in demyelinated (csn) zebrafish treated with Y27632 ROCK inhibitor. Arrows indicate the position of the cell body. Arrowheads indicate areas of mistargeted myelin. Scale bars, 20 µm. (E) Fate of the 23 oligodendrocytes that survived demyelination: 9 cells formed new sheaths only, 6 cells formed new sheaths and mistargeted myelin, 7 cells formed mistargeted myelin only, and 1 cell formed no new myelin. (F) Quantification of the number of mistargeted myelin profiles produced by the same oligodendrocytes pre-treatment (median = 0.00, 25^th^ percentile 0.00, 75^th^ percentile 0.00) and 3 dpt (median = 1.00, 25^th^ percentile 0.00, 75^th^ percentile 1.00), p = 0.0012. Wilcoxon test. N = 23 oligodendrocytes from 23 zebrafish (pre-treatment and surviving). (G) Quantification of the number of myelin sheaths produced by the same oligodendrocytes pretreatment (median = 18.00, 25^th^ percentile 13.00, 75^th^ percentile 20.00) and 3 dpt (median = 2.00, 25^th^ percentile 1.00, 75^th^ percentile 3.00), p < 0.0001. Wilcoxon test. N = 23 oligodendrocytes from 23 zebrafish (pre-treatment and surviving). (H) Quantification of the average length of myelin sheaths produced by the same oligodendrocytes pre-treatment (mean = 30.93 ± 5.76 SD) and 3 dpt (mean = 25.47 ± 9.16 SD), p = 0.0655. Paired two-tailed t-test. N = 23 oligodendrocytes from 23 zebrafish (pre-treatment and surviving). (I) Quantification of the number of mistargeted myelin profiles per oligodendrocyte following demyelination (of the oligodendrocytes that mistargeted myelin) at 3 dpt in DMSO (mean = 1.40 ± 0.89 SD) versus Y27632 ROCK inhibitor (mean = 3.50 ± 2.27 SD), p = 0.0414. Welch’s t-test. N = 5 - 8 zebrafish per condition. (J) Quantification of the average number of sheaths produced per oligodendrocyte at 3 dpt in control DMSO (median = 3.00, 25^th^ percentile 1.50, 75^th^ percentile 3.00) versus Y27632 ROCK inhibitor (median = 4.00, 25^th^ percentile 1.00, 75^th^ percentile 4.50), p = 0.5964. Mann-Whitney test. N = 5 - 9 zebrafish per condition.

### Newly generated oligodendrocytes have a large capacity for remyelination and rarely mistarget myelin

To directly compare remyelination by surviving oligodendrocytes and newly generated cells, we next investigated the remyelination potential of new oligodendrocytes generated after demyelination (**Figure 4, Methods**). As indicated by our global analyses of oligodendrocytes and myelination (**Figure 2B+C**), we found that newly generated oligodendrocytes and their myelin sheaths appeared along the same tracts as controls, i.e. those that are myelinated at this age. We found that newly generated oligodendrocytes exhibit a very high remyelination capacity and made an average of 25 sheaths per cell (**Figure 4A,C**), almost all of which were targeted to axons (509 correctly targeted sheaths versus 7 mistargeted myelin profiles in 20 oligodendrocytes) (**Figure 4B**). This indicates that previously demyelinated axons are permissive for remyelination, and that the remyelination capacity of new cells is more extensive than that of surviving oligodendrocytes, which we studied over the same period of time and in the same region. We noted that newly generated oligodendrocytes had more sheaths than oligodendrocytes had prior to demyelination (compare **Figure 4C** with **Figure 3G**), but that the average length of these sheaths was shorter (compare **Figure 4D** with **Figure 3H**). Although short myelin sheaths have long been considered a hallmark of remyelination (42), we asked whether this might simply be due to the production of more sheaths per cell. To address this, we estimated the total myelinating capacity of oligodendrocytes (by multiplying sheath number by average length) and found no significant difference between newly generated oligodendrocytes and oligodendrocytes before demyelination (**Figure 4E**). This suggests that the short myelin sheaths of newly generated oligodendrocytes may reflect differences in the distribution of the myelin produced by these cells over time, and that short sheaths may eventually grow to a normal size, as has been demonstrated in other models (43, 44). Our analyses of total myelin production also showed that in comparison to oligodendrocytes that survive demyelination, newly generated cells make over 10-fold more myelin (**Figure 4E**), indicating that their potential for remyelination is far greater than that of oligodendrocytes that survive demyelination.

**Fig. 4.**
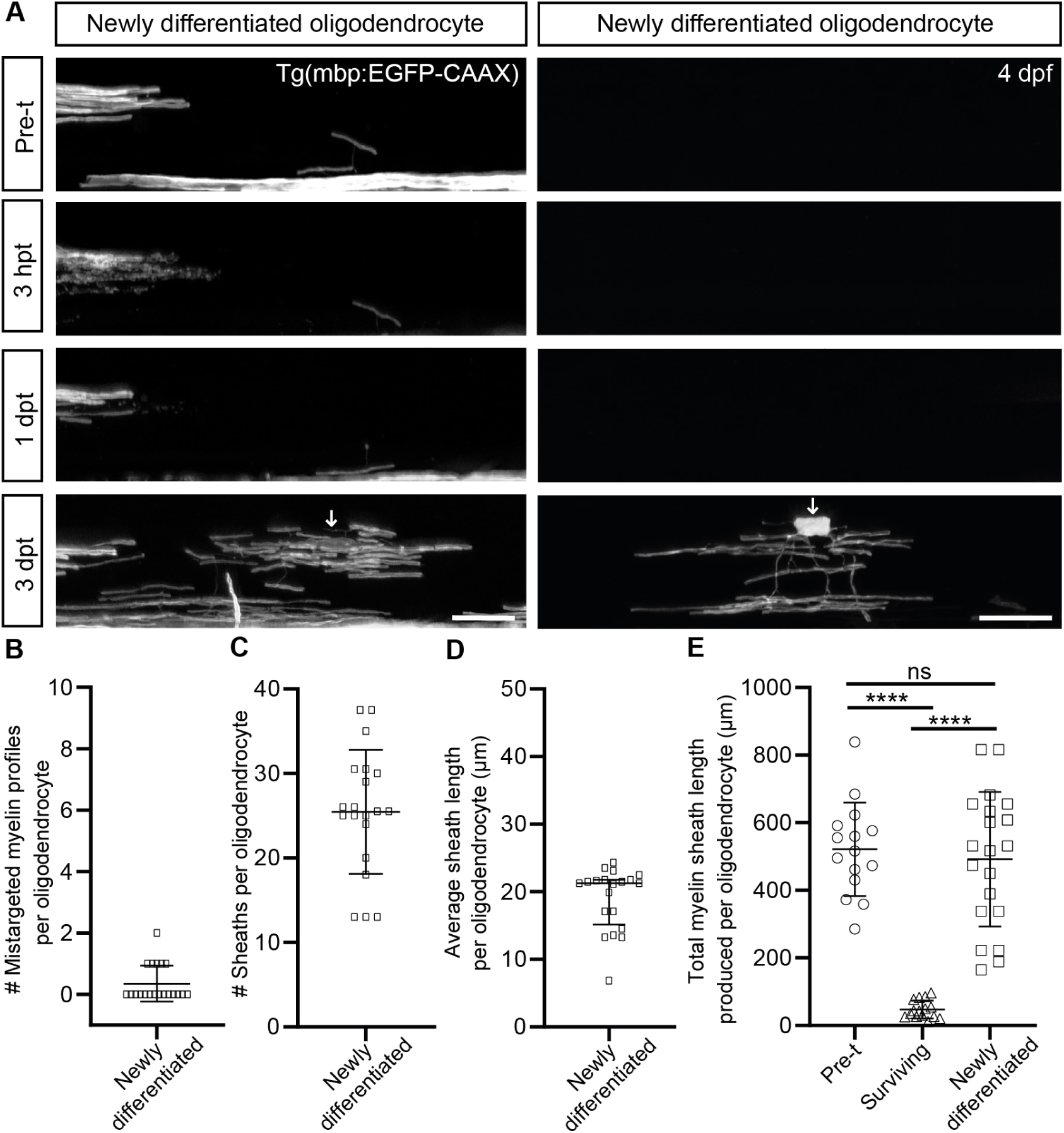
Extensive remyelination by newly generated oligodendrocytes. (A) Confocal images of csn treated zebrafish with oligodendrocytes generated after demyelination. Arrows show position of cell bodies. Scale bars, 20 µm. (B) Quantification of the average number of mistargeted myelin profiles produced per newly generated oligodendrocyte at 3 dpt (median = 0.00, 25^th^ percentile 0.00, 75^th^ percentile 1.00). N = 20 oligodendrocytes from 11 zebrafish. (C) Quantification of the average number of myelin sheaths produced per newly generated oligodendrocyte at 3 dpt (mean = 25.45 ± 7.33 SD). N = 20 oligodendrocytes from 11 zebrafish. (D) Quantification of the average length of myelin sheaths produced per newly generated oligodendrocyte at 3 dpt (median = 21.25, 25^th^ percentile 15.17, 75^th^ percentile 21.76). N = 20 oligodendrocytes from 11 zebrafish. (E) Quantification of the total myelin produced per oligodendrocyte (calculated by multiplying number of sheaths per oligodendrocyte by the average sheath length per oligodendrocyte) pre-treatment (mean = 521.5 ± 138.30 SD), versus the same oligodendrocytes 3 dpt (mean = 47.18 ± 26.57 SD) and by newly differentiated oligodendrocytes at 3 dpt (mean = 491.80 ± 199.10 SD). Pre-treatment versus surviving p < 0.0001, pre-treatment versus newly differentiated p = 0.8271, surviving versus newly differentiated p < 0.0001. Ordinary one-way ANOVA with Tukey multiple comparison test. N = 15 oligodendrocytes from 15 zebrafish (pre-treatment and surviving). N = 20 oligodendrocytes from 11 zebrafish (newly differentiated).

## Discussion

Our study aimed to combine analyses of oligodendrocytes in MS with investigation of the dynamic nature of remyelination in a transgenic zebrafish line that allowed longitudinal *in vivo* imaging of individual oligodendrocytes. We designed a model that allowed us to assess remyelination by both newly generated oligodendrocytes, that have long been known to have regenerative capacity, and by oligodendrocytes that survive demyelination, that have more recently been implicated in remyelination, including in MS. Our analyses revealed a new feature of oligodendrocytes in MS, the mistargeting of myelin to neuronal cell bodies, which we found was also a characteristic of remyelination by surviving oligodendrocytes in zebrafish. In addition to myelin mistargeting, we found that surviving oligodendrocytes exhibited a very limited capacity to form new sheaths. Nonetheless, when sheaths were made by surviving oligodendrocytes, their growth along axons appeared relatively normal, indicating that surviving cells retain the capacity to support sheath elongation. In contrast, our analyses revealed robust remyelination by newly generated oligodendrocytes, which made an even greater number of sheaths than oligodendrocytes had prior to demyelination. Our data provide insights into the limited remyelination potential of surviving oligodendrocytes, and suggest that newly generated cells may have a greater capacity to promote remyelination in conditions like MS (see **Summary schematic**).

Our observations of oligodendrocytes that survive demyelination indicate that these cells do not have a robust capacity to reinstate a programme of dynamic process extension, target selection or sheath formation. This is in line with recent evidence from studies of remyelination in the rodent cortex, where surviving oligodendrocytes were shown to have the capacity to generate new sheaths, but made very few per cell (14). This reflects previous observations which indicate that myelin sheath formation occurs rapidly over a short period (26). Given that surviving oligodendrocytes exhibited such a limited capacity for sheath formation, we tried to enhance their process extension by treating animals with inhibitors of ROCK, but this served only to further the mistargeting of myelin. It remains to be determined if surviving oligodendrocytes can be manipulated at all to exhibit truly robust sheath formation. One possibility would be to reprogramme surviving oligodendrocytes to a less mature state from which they can reinstate dynamic process extension, target selection and sheath formation. Our study did, however, indicate that surviving oligodendrocytes can support the elongation of sheaths along axons. This observation also fits with the fact that mature oligodendrocytes exhibit a protracted, potentially life-long ability to remodel existing myelin sheaths, including in humans (45). Therefore, it is possible that surviving oligodendrocytes may be best poised to remyelinate axons close to their cell body, or where oligodendrocyte processes remain associated with axons after demyelination, or in situations where myelin sheaths sustain only partial damage. Indeed, a recent study has indicated that neural circuit activity can promote remyelination by surviving oligodendrocytes (14). Furthermore, it might be possible to enhance the remyelination capacity of surviving oligodendrocytes by manipulating specific signalling pathways that can drive the growth of mature sheaths (27, 28).

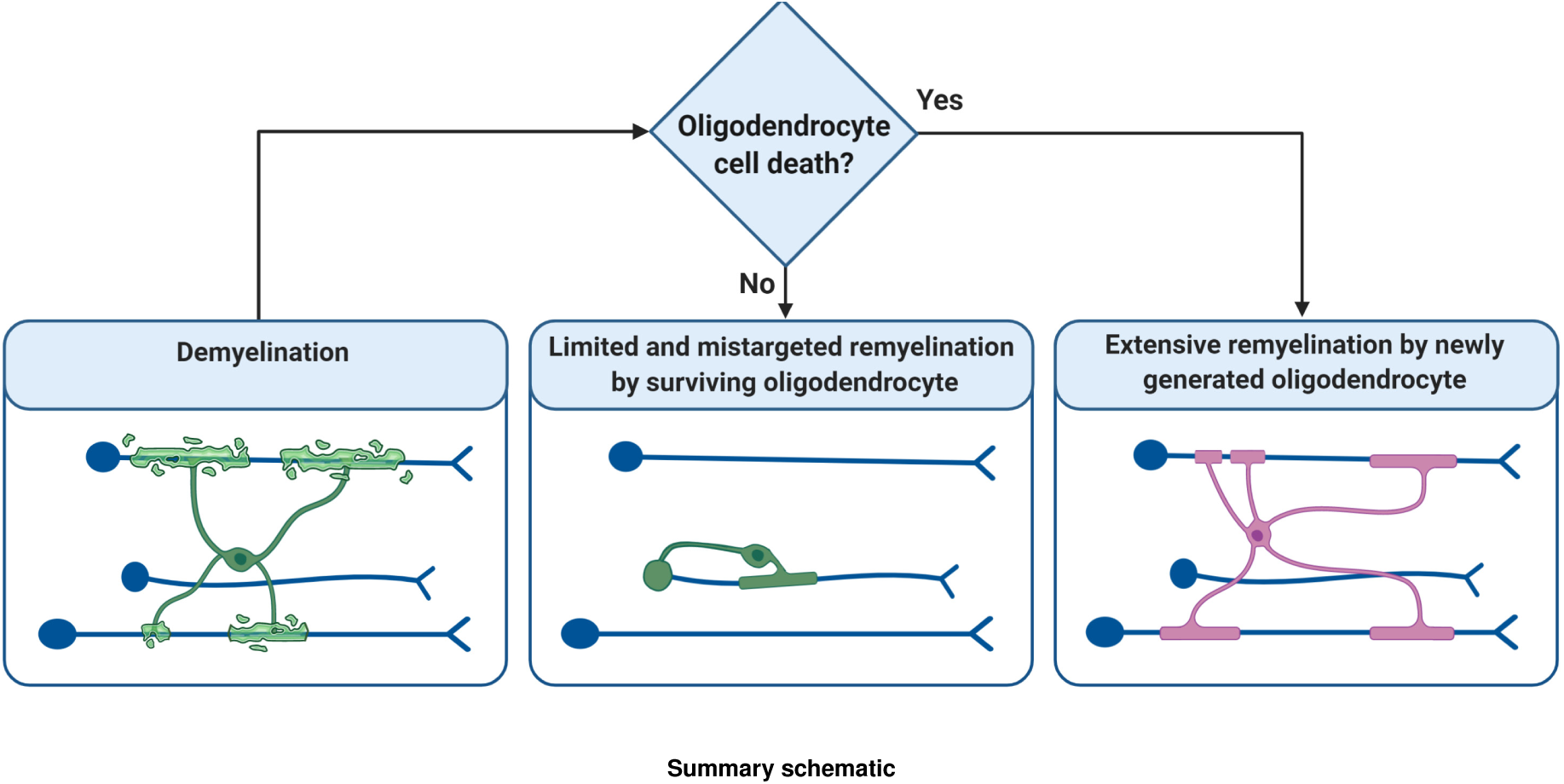
Summary schematic

Our study directly compared the regenerative capacity of newly generated and surviving oligodendrocytes *in vivo*, indicating that the regenerative potential of newly generated oligodendrocytes is superior to that of surviving cells. Newly generated cells made an abundant number of new sheaths, rarely mistargeted myelin, and had a total capacity for remyelination ten-fold higher than that of survivors. To date, therapeutic strategies to promote remyelination have focussed on increasing the generation of new oligodendrocytes (3), and our data indicate that this remains a promising approach. Remyelination in MS is known to vary between individuals, and it has recently been shown that some people with MS exhibit higher levels of new oligodendrocyte generation than others (13). It will be important for future studies to determine whether the relative number of newly made oligodendrocytes compared to surviving oligodendrocytes relates to remyelination efficiency in people with MS. The presence of surviving oligodendrocytes may even impair remyelination. It has been shown in animal models, both in the zebrafish spinal cord (46) and mammalian cortex (47), that oligodendrocyte cell death leads to a homeostatic generation of new cells. Therefore, it is possible that the presence of oligodendrocytes that survive demyelination means that the signals associated with cell death that trigger OPCs to produce new oligodendrocytes are absent. If so, it may even be worth considering targeted destruction of surviving cells, to allow activation of OPCs and the generation of new oligodendrocytes with better remyelinating potential, a possibility that will require investigation in experimental models. In summary, our analyses reveal a striking new pathology in MS, that may reflect limited and aberrant remyelination by oligodendrocytes that survive demyelination. Our study also indicates that targeting the generation of new oligodendrocytes, and their capacity for remyelination, may represent the best strategy to promote myelin regeneration for the treatment demyelinating disorders such as MS.

## Methods

### Human data

Post-mortem brain tissue (motor cortices) from MS patients and controls without neurological defects were provided by a UK prospective donor scheme with full ethical approval from the UK Multiple Sclerosis Society Tissue Bank (MREC/02/2/39). MS diagnosis was confirmed by neuropathological means by F. Roncaroli (Imperial College London) and clinical history was provided by R. Nicholas (Imperial College London). **Table 1** includes details on samples used. Tissue blocks of 2 cm x 2 cm x 1 cm were collected, fixed, dehydrated and embedded in paraffin blocks. 4 µm sequential sections were cut and stored at room temperature. Grey matter MS lesions were identified using anti-Proteolipid Protein (PLP) immunostaining.

**Table 1.** Information on human donor tissue. A list of human donor tissue with information on sex (M=male, F=female), age at death, cause of death (COPD=chronic obstructive pulmonary disease, SPMS=secondary progressive multiple sclerosis), post-mortem interval and MS disease duration.

| Sample ID | Sex | Age (years) | Cause of death | Post-mortem interval (hours) | MS disease duration (years) |
| --- | --- | --- | --- | --- | --- |
| CO72 | M | 77 | Pneumonia, ischaemic bowel | 26 | N/A |
| CO73 | M | 71 | Liver cancer | 29 | N/A |
| CO74 | F | 84 | Old age | 22 | N/A |
| CO75 | M | 88 | COPD | 8 | N/A |
| CO76 | M | 87 | Pneumonia, Idiopathic pulmonary fibrosis | 31 | N/A |
| MS239 | F | 69 | Bronchopneumonia, SPMS | 42 | 51 |
| MS243 | M | 61 | Sigmoid volvulus, SPMS | 77 | 40 |
| MS250 | F | 60 | Bronchopneumonia, SPMS | 38 | 32 |
| MS285 | M | 61 | Paracetamol overdose, SPMS | 108 | 21 |
| MS510 | F | 38 | Pneumonia, SPMS | 31 | 22 |

### Human post-mortem brain tissue immunohistochemistry

Paraffin sections were rehydrated, washed in PBS and microwaved for 15 minutes in Vector Unmasking Solution for antigen retrieval (H-3300, Vector). For colorimetric immunohistochemistry, endogenous peroxidase and alkaline phosphatase activities were blocked for 10 minutes using Bioxal solution (SP-6000, Vector). Sections were then blocked with % normal horse serum (S-2012, Vector) for 1 hour at room temperature. Primary antibodies were incubated in antibody diluent solution (003118, Thermo Fisher Scientific), overnight at 4 °C in a humidified chamber. Horse peroxidase or alkaline phosphatase-conjugated secondary antibodies (Vector) were applied for an hour at room temperature. Staining development was performed using either DAB HRP substrate kit or Vector Blue substrate kit (both from Vector) according to the manufacturer’s instructions.

### Human post-mortem brain tissue immunofluorescence

For immunofluorescence, sections were incubated with Autofluorescent Eliminator Reagent (2160, MERCK-Millipore) for 1 minute and briefly washed in 70 % ethanol after antigen retrieval. The sections were subsequently incubated with Image-iT^®^ FX Signal Enhancer (I36933, Thermo Fisher Scientific) for 30 minutes at room temperature, washed and blocked for 1 hour with 10 % normal horse serum, 0.3 % Triton-X in PBS. Primary antibodies were diluted in antibody diluent solution (as above) and placed on sections overnight at 4 °C in a humidified chamber. The next day the sections were incubated with Alexa Fluor secondary antibodies (Thermo Fischer Scientific, 1:1000) for 1 hour at room temperature and counterstained with Hoechst for the visualization of the nuclei. Primary antibodies used: rabbit polyclonal IgG antibody to NeuN (104225, Abcam, 1:100) and mouse monoclonal IgG2A antibody to myelin Proteolipid Protein (clone PLPC1, MAB388, MERCK-Millipore, 1:100). All slides were mounted using Mowiol mounting medium (475904, MERCK-Millipore).

Entire sections were imaged using the Zeiss AxioScan Slide scanner and all quantifications were performed using Zeiss Zen lite imaging software. For all cases, the whole grey matter area of the section was investigated focusing on perilesion areas in MS cases and in corresponding cortical layers in control tissue. Cell densities are presented as immune-positive cells per cm^2^. Z-stack images of the fluorescent-labelled samples were acquired with the LSM 880 confocal microscope equipped with Airyscanner and a 20X objective (Zeiss Plan-Apochromat 20X dry, NA = 0.8). Z-stacks were acquired with an optimal z-step according to the experiment.

### Zebrafish lines and maintenance

All zebrafish were maintained under standard conditions in the Queen’s Medical Research Institute CBS Aquatics facility at the University of Edinburgh. Studies were carried out with approval from the UK Home Office and according to its regulations, under project licenses 70/8436 and PP5258250. Adult animals were kept in a 14 hours light and 10 hours dark cycle. Embryos were kept at 28.5 °C in 10 mM HEPES-buffered E3 embryo medium or conditioned aquarium water with methylene blue. Larval zebrafish were analysed between 4 - 9 dpf, before the onset of sexual differentiation in zebrafish.

The following transgenic zebrafish (Danio rerio) lines were used in this study: Tg(mbp:TRPV1-tagRFPt) (generated for this study, see details below), Tg(mbp:EGFP) and Tg(mbp:EGFP-CAAX) (36). Throughout the text and figures ‘Tg’ denotes stable, germline inserted transgenic line.

### Generation of the Tg(mbp:TRPV1-tagRFPt) line

We reasoned that we might be able to induce primary disruption to myelin sheaths by generating a transgenic system in which we drove cationic influx into myelin. To do so, we generated the Tg(mbp:TRPV1-tagRFPt) transgenic line, in which the cation-permeable TRPV1 channel of rats was expressed in myelinating oligodendrocytes downstream of the myelin basic protein (mbp) promoter of zebrafish. Zebrafish TRPV1 is not sensitive to capsaicin, meaning that only cells expressing the rat form of TRPV1 are capsaicin sensitive and it is only in those cells that capsaicin-induced cation influx takes place: the principle of this system has previously been used to selectively ablate neurons in zebrafish (35). The mbp:TRPV1-tagRFPt plasmid was generated by first making a pME-TRPV1-tagRFPt entry vector plasmid. To do this, the TRPV1-tagRFPt sequence was amplified by PCR from the (islet1:GAL4VP16,4xUAS:TRPV1-RFPT) plasmid used by (35), using the primers in **Table 2**. This PCR fragment was then inserted into the backbone pDONR221 by a BP reaction (BP Clonase II, Thermo Fisher Scientific). The entry vectors p5E-mbp-promoter, pME-TRPV1-tagRFPt, p3E-polyA, and pDest-Tol2pA2 were recombined by an LR reaction (LR Clonase II, Thermo Fisher Scientific) using the multisite Gateway system (48) to generate the final plasmid. Finally, 1-2 nl of 10 - 15 ng/µl plasmid DNA was injected into wild-type eggs along with 25 ng/µl tol2 transposase mRNA. Injected larvae were screened for mosaic transgene expression, and transgene positive larvae raised to adulthood (F0 generation). F0 generation zebrafish were then outcrossed and their progeny (F1 generation) screened for full expression of the transgene.

**Table 2.**
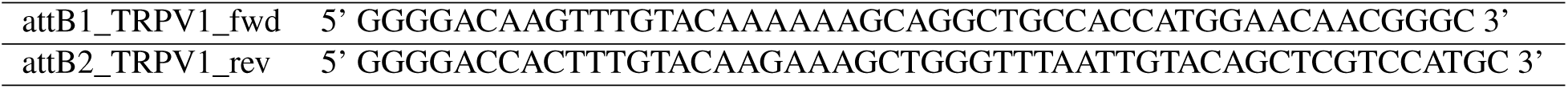
Primers used to generate the Tg(mbp:TRPV1-tagRFPt) line.

### Capsaicin induction of demyelination

Capsaicin (csn) (Sigma-Aldrich) was prepared as a 20 mM primary stock in 100 % DMSO (Fisher Scientific) and stored at −80 °C. Concentrations of csn were titrated to find an optimal dose to induce severe demyelination without affecting overall zebrafish health. 10 µM csn 1 % DMSO was selected as the lowest dose to induce severe demyelination in a 2 hour treatment period without inducing any visible behavioural response, paralleling previous results (35). We further tested the specificity of the 10 µM csn 1 % DMSO 2 hour treatment on zebrafish which did not express rat TRPV1 channels (**Figure S2**). We observed no effect on oligodendrocytes or the myelin they produced in zebrafish which did not contain the Tg(mbp:TRPV1-tagRFPt) transgene.

Treatments of 10 µM csn in 1 % DMSO or 1 % DMSO alone (vehicle control referred to as DMSO in figures) in E3 medium were applied by bath application to larval zebrafish for 2 hours at 28.5 °C. Following treatment, to ensure no capsaicin or DMSO remained all solutions were replaced three times with fresh E3 embryo medium.

### Y27632 treatment

Y27632 (Tocris) was prepared as a 200 mM primary stock in 100 % DMSO and stored at −80 °C. Zebrafish were demyelinated with capsaicin treatment as previously described at 4 dpf. Immediately after washing out the csn treatment, 100 µM Y27632 in 1 % DMSO or 1 % DMSO alone (vehicle control referred to as DMSO in figures) in E3 medium were applied by bath application to larval zebrafish between 4 dpf (3 hpt) to 7 dpf (3 dpt) with daily media changes. Zebrafish were kept at 28.5 °C between imaging timepoints.

### Transmission electron microscopy (TEM)

Tissue was prepared for TEM as previously described (49). Briefly, zebrafish embryos were terminally anaesthetised in MS222 tricaine (Sigma-Aldrich) and incubated, with microwave stimulation, first in primary fixative (4 % paraformaldehyde + 2 % glutaraldehyde in 0.1 M sodium cacodylate buffer) and then in secondary fixative (2 % osmium tetroxide in 0.1 M sodium cacodylate/imidazole buffer). Samples were then stained en bloc with saturated uranyl acetate solution and dehydrated in an ethanol and acetone series, both with microwave stimulation. Samples were embedded in EMbed-812 resin (Electron Microscopy Sciences) and sectioned using a Reichert Jung Ultracut Microtome. Sections were cut at comparable somite levels by inspection of blocks under a dissection microscope and stained in uranyl acetate and Sato lead stain. TEM images were taken with a Phillips CM120 Biotwin TEM. The Photomerge tool in Adobe Photoshop was used to automate image registration and tiling. To assess axon diameter (> 0.4 µm), axonal areas were measured in ImageJ.

### Single oligodendrocyte labelling

To mosaically label oligodendrocytes, fertilized eggs were injected with 1 nl of 10 ng/µl pTol2-mbp:EGFP-CAAX plasmid DNA and 50 ng/µl tol2 transposase mRNA at the 1 cell stage. Animals were screened at 3 and 4 dpf for isolated oligodendrocytes.

### Live imaging

Larval zebrafish were live imaged after anesthetising them with MS222 tricaine and mounting them on glass coverslips, embedded in 1.5 % low melting point agarose (Invitrogen). Z-stacks were acquired at relevant locations along the spinal cord using the LSM 880 confocal microscope equipped with Airyscan Fast and a 20X objective (Zeiss Plan-Apochromat 20X dry, NA = 0.8). Z-stacks were acquired with an optimal z-step according to the experiment. To follow the fate of oligodendrocytes, and myelination profiles over time, zebrafish were repeat imaged (4 dpf pretreatment, 3 hpt, 1 dpt, and 3 dpt). To do so zebrafish were carefully cut out from agarose following imaging and returned to E3 embryo medium, whilst being checked for signs of impaired health or swim behaviour. Zebrafish were maintained at 28.5 °C in 10 mM HEPES-buffered E3 embryo medium between imaging sessions.

To assess any developmental effects of the Tg(mbp:TRPV1-tagRFPt) transgene zebrafish were anesthetised with MS222 tricaine and imaged using the Vertebrate Automated Screening Technology BioImager system (Union Biometrica), which automated the positioning of larvae and image acquisition using a Zeiss spinning disk confocal microscope (VAST-SDCM) (50).

### Single oligodendrocyte imaging

Zebrafish were screened at 4 dpf for isolated oligodendrocytes, and when identified were imaged over time by identifying their position along the spinal cord, relative to the nearest somite as well as their position relative to other oligodendrocytes in the surrounding area labelled with the Tg(mbp:TRPV1-tagRFPt) and Tg(mbp:EGFP-CAAX) transgenes. As not all oligodendrocytes die after initial demyelination, we were able to carefully assess relative positions using both tagRFPt and EGFP-CAAX reporters (**Figure S2E**).

### Image analysis

Image processing and analysis was performed in Fiji (ImageJ). Figure panels were produced using Fiji and Adobe Illustrator. For figures, maximum-intensity projections of z-stacks were made, and a representative x-y area was cropped. All zebrafish images represent a lateral view of the spinal cord, anterior to the left and dorsal on top. For most images, processing included only global change of brightness and contrast; further processing and analysis is as follows.

### Cell counts

Mbp:EGFP cell bodies filled with cytoplasmic EGFP were counted through z-stacks which encompassed the depth of the spinal cord using the cell counter plugin in Fiji. The same area of the spinal cord was imaged and counted for analysis in different zebrafish (5 somite section). Dorsal and ventral oligodendrocytes were counted separately and combined for total analysis. All images were blinded to treatment condition during analysis.

### Single oligodendrocyte morphology analysis

Oligodendrocytes were analysed for sheath length and number using the segmented line tracing tool in Fiji (Image J) through z-stacks which encompassed the depth of the oligodendrocyte and its myelin. Regions of interest were saved per oligodendrocyte. Cell body counts were carried out manually using the full z-stack of each oligodendrocyte. Oligodendrocytes with too much overlapping myelin, from oligodendrocytes in the surrounding area, to confidently trace the source of the myelin sheath were excluded from analysis.

### Reproducibility

Larval zebrafish were grown up in the same incubator to avoid any differences due to developmental stage between experiments. Following screening to identify transgene positive zebrafish, larvae were randomly assigned to treatment or control groups. During live imaging analyses, experimental and control animals were imaged in an alternating pattern (per experiment) to ensure no confounding effects of developmental stage between groups.

### Statistics and reproducibility

Graphs and statistical tests were carried out using GraphPad Prism. Analysis of *in vivo* data was carried out blinded before treatment group was revealed. Data were averaged per biological replicate (N represents number of zebrafish or humans) unless stated otherwise in figure legends.

Data was tested for normal distribution using the D’Agostino & Pearson test or the Shapiro-Wilk test. The variance of the data was assessed using the F test for variance. Indicated p values are from two-tailed unpaired t-tests, two-tailed unpaired Welch’s t-tests, two-tailed paired t-tests, Wilcoxon signed-rank tests, or Mann-Whitney tests. To compare more than 2 groups a one-way ANOVA with Tukey’s multiple comparisons test or a Kruskal-Wallis test with Dunn’s multiple comparisons test was used. A difference was considered statistically significant when p < 0.05. Throughout the figures p values are indicated as follows: not significant or ‘ns’ p > 0.05, ‘*’ p < 0.05, ‘**’ p < 0.01, ‘***’p < 0.001. All data are shown as mean ± standard deviation (SD) where data is normally distributed or median with interquartile range (25^th^ percentile and 75^th^ percentile) where data is not normally distributed. Details of statistical test used, precise p values and N values for each comparison are detailed in figure legends.

### Illustrations/Schematics

Illustrations were created with Biorender or Adobe Illustrator.

## Acknowledgements

We thank Charles ffrench-Constant, Ethan Hughes and members of the Lyons lab for critical feedback, the BVS aquatics facility for fish care, and Carmen Melendez-Vasquez for suggesting the ROCK inhibitor experiment. This work was supported by Wellcome Trust Senior Research Fellowships (102836/Z/13/Z and 214244/Z/18/Z), a Medical Research Council Project Grant (MR/P006272/1) and an MS Society Innovative Grant (95) to DAL. SAN and JMW were supported by a Wellcome Trust Four-Year Ph.D. Program in Tissue Repair (Grant 108906/Z/15/Z) and JMW also by a University of Edinburgh Ph.D. Tissue Repair Studentship Award (MRC Doctoral Training Partnership MR/K501293/1). LZ and AW were supported by MS Society UK Centre grant.

## Author Contributions

SAN, JMW and DAL conceived the project. SAN, JMW, AK and JJE designed and performed the *in vivo* experiments. LZ and AW designed and performed the human tissue experiments. SAN and DAL co-wrote the manuscript, edited by all. DAL managed the project.

## Supplementary Figures

**Fig. S1.**
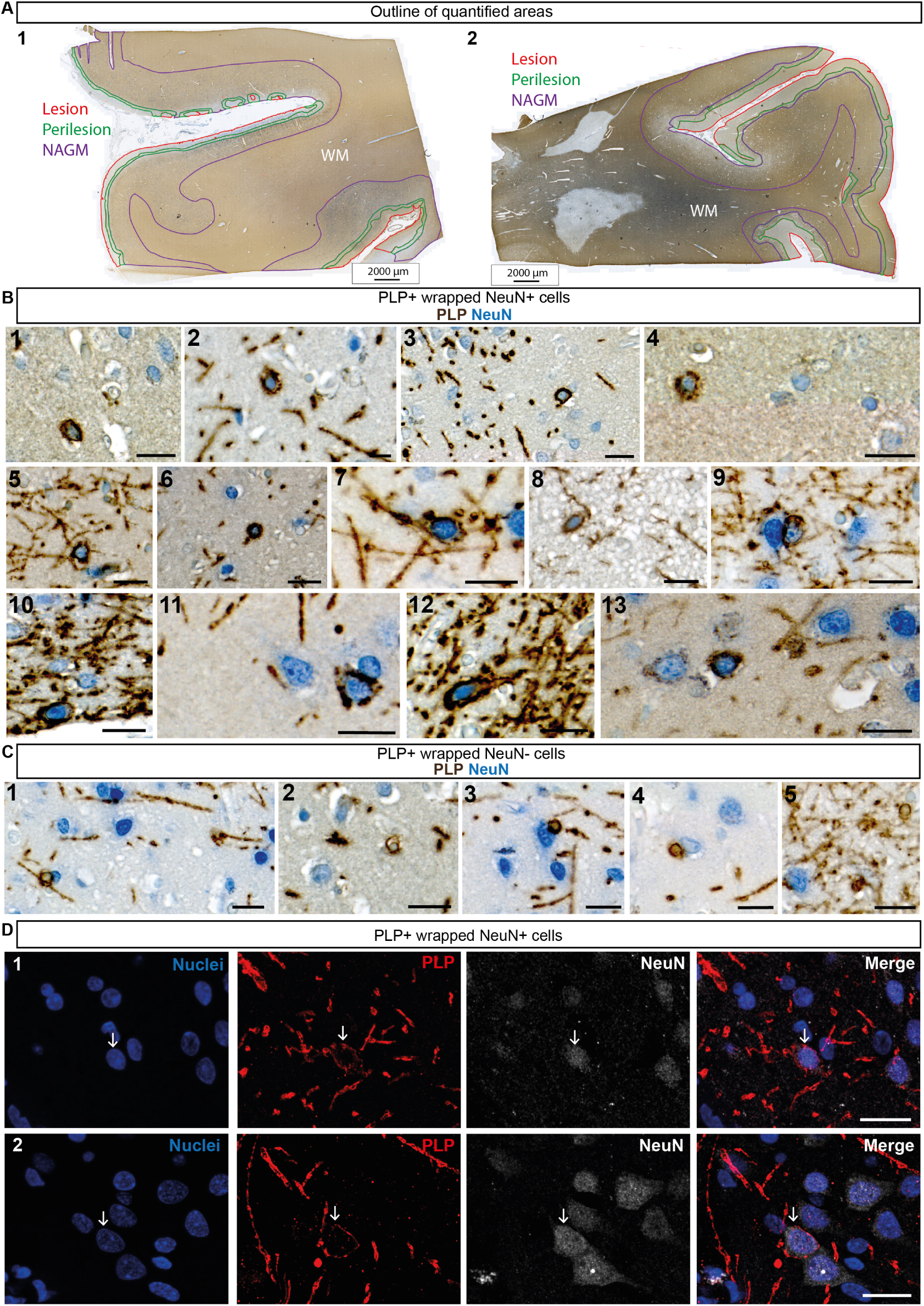
Mistargeted myelin profiles are present in remyelinating lesions in motor cortex tissue from people with MS. (A) Low magnification image of chromogenic immunohistochemistry for proteolipid protein (PLP - brown) and NeuN (blue) in human MS motor cortex. Outline of quantified areas shown with lesion area highlighted in red, perilesion area highlighted in green, normal appearing grey matter (NAGM) in purple and white matter indicated by ‘WM’ in white. Images 1 and 2 show examples of quantified areas in 2 different human MS motor cortex samples. Scale bars, 2000 µm. (B and C) High magnification images of chromogenic immunohistochemistry for proteolipid protein (PLP - brown) and NeuN (blue) in human MS motor cortex. (B) Images 1 - 13 show example images of PLP +ve wrapped NeuN +ve cells (myelinated neuronal cell bodies). Images 1 - 13 scale bars, 20 µm. Image 2 scale bar, 10 µm. (C) Images 1 - 5 show example images of PLP +ve wrapped NeuN -ve cells (oligodendrocytes). Scale bars, 20 µm. (D) Fluorescent immunohistochemistry for NeuN (white), PLP (red) and Hoechst (nuclei - blue) in human MS motor cortex. Arrow indicates location of PLP +ve wrapped NeuN +ve Hoechst +ve cells (myelinated neuronal cell bodies). Images 1 and 2 represent different PLP +ve wrapped NeuN +ve Hoechst +ve cells. Scale bars, 20 µm.

**Fig. S2.**
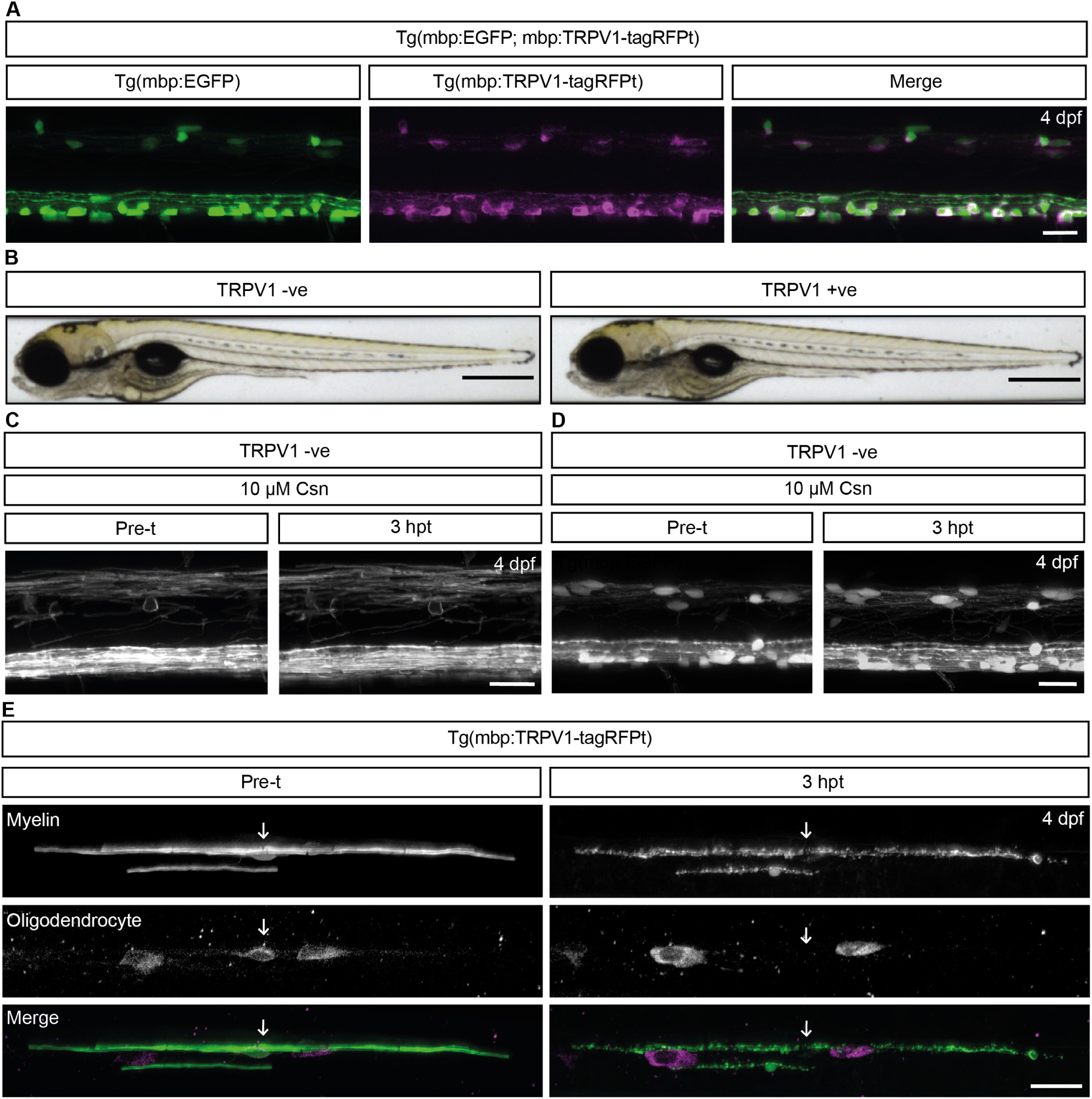
Characterisation of the Tg(mbp:TRPV1-tagRFPt) demyelination line. (A) Confocal images of myelinating oligodendrocytes in the Tg(mbp:EGFP; mbp:TRPV1-tagRFPt) zebrafish line at 4 dpf showing overlap in the EGFP and tagRFPt channels in the merged image. Scale bar, 20 µm. (B) Brightfield images of a zebrafish containing the Tg(mbp:TRPV1-tagRFPt) transgene (TRPV1 +ve), or wildtype siblings without the Tg(mbp:TRPV1-tagRFPt) transgene (TRPV1 -ve) at 4 dpf. Scale bar = 500 µm. (C and D) Confocal images of the (C) Tg(mbp:EGFP-CAAX) line and the (D) Tg(mbp:EGFP) line pre-treatment (indicated here as pre-t) at 4 dpf, and 3 hpt. Zebrafish not containing the Tg(mbp:TRPV1-tagRFPt) transgene show no disruption to myelin or oligodendrocytes following a 2 hour treatment of 10 µM csn. Scale bars, 20 µm. (E) Confocal images of a single oligodendrocyte labelled with mbp:EGFP-CAAX in the Tg(mbp:TRPV1-tagRFPt) line pre-treatment (pre-t) at 4 dpf, and 3 hpt. An example of oligodendrocyte cell death is demonstrated here by the disappearance of a tagRFPt +ve cell body following csn treatment in the same zebrafish before and after demyelination. Arrows indicate the location of the cell body, or where it was prior to demyelination. Scale bar, 20 µm.

